# Cortical states control visual spatial perception

**DOI:** 10.1101/316398

**Authors:** Anderson Speed, Joseph Del Rosario, Christopher P. Burgess, Bilal Haider

## Abstract

Many factors modulate the state of cortical activity, but the importance of cortical states for sensory perception remains debated. We trained mice to detect spatially localized visual stimuli, and simultaneously measured local field potentials and excitatory and inhibitory neuron populations across layers of primary visual cortex (V1). Cortical states with low firing rates and correlations between excitatory neurons, and reduced oscillatory activity in Layer 4, accurately predicted single trials of visual spatial detection behavior. Our results show that cortical states exert strong effects at the initial stage of cortical processing in V1, and play a decisive role for visual spatial behavior in mice.

## Introduction

Behavioral factors such as sleep, wakefulness, and movement have strong effects on the state of cortical activity. Cortical states are typically defined by the degree of shared fluctuations among cortical neural populations, measured by local field potential (LFP) frequency power (Harris and Thiele, 2011), and neural population correlations (Kohn et al., 2009). Cortical states exert profound effects on sensory responses (Haider and McCormick, 2009; Petersen and Crochet, 2013), but there remain unresolved questions about cortical states and their effects on sensory perception.

One question concerns the role of cortical states for perception across sensory modalities. In primates performing visual tasks, cortical states strongly influence stimulus detection (Gilbert and Li, 2013; Reynolds and Chelazzi, 2004; Spitzer et al., 1988). However, recent studies in mice show that somatosensory perception occurs in a wide variety of cortical states—even those with correlations and activity patterns similar to sleep (Sachidhanandam et al., 2013). It is unknown how cortical states influence visual perception in mice, and if the underlying mechanisms are like those in other mammalian visual systems, or like those in other mouse sensory cortical areas.

A second question concerns how cortical states coordinate excitatory and inhibitory neuron population activity during perception. In primates performing visual tasks, selective attention strongly modulates cortical state (Engel et al., 2016) and population correlations (Kohn et al., 2016; Nienborg et al., 2012). Reduction of correlated activity (decorrelation) best accounts for perceptual improvements in these tasks (Cohen and Maunsell, 2009), but the neuronal subtypes involved remain unclear. Identifying cortical neuron subtypes in higher mammals presents challenges, since action potentials of many excitatory neurons are indistinguishable from those of inhibitory neurons (Constantinople et al., 2009; Haider et al., 2010; Soares et al., 2017; Vigneswaran et al., 2011). This may hinder full understanding of excitatory and inhibitory contributions to decorrelation and sensory perception.

A third question concerns how cortical states affect information flow across cortical layers during sensory perception. A recent study of primate visual cortex area V4 revealed that cortical states underlying selective visual attention strongly modulate correlations in the input layers (Nandy et al., 2017). In contrast, input layers in primary visual cortex (V1) exhibit low correlations and low sensitivity to cortical state changes (Hansen et al., 2012; Poort et al., 2016; Smith et al., 2013). It remains unknown how cortical states modulate activity across input and output layers of V1 during visual perception in mice.

To address these unresolved questions, we trained mice to detect visual stimuli appearing in discrete portions of the visual field, and simultaneously measured LFP and excitatory and inhibitory neuron populations across layers of V1. We found that cortical states with low excitatory neuron firing rates, decorrelated populations, and suppressed LFP oscillations accurately predicted stimulus detection.

## Results

### Visual detection latency and sensitivity depend upon spatial location

We designed a behavioral assay of visual spatial perception in stationary head-fixed mice (Fig. 1A). Mice reported detection of visual stimuli by licking for water rewards. These were obtained only if they licked during the stimulus window (typically 1 – 1.5 s). Stimuli appeared only after a mandatory period of no licks had elapsed (typically 0.5 – 6 s, randomized per trial), and stimuli disappeared upon the first lick during the stimulus window. Static horizontally oriented Gabor gratings appeared in one of two fixed spatial locations, either in the monocular or binocular visual fields. Gratings appeared at one of these locations for a block of 15 to 50 consecutive trials, and then switched to the other location for a new block of trials (Fig. 1B). During training, we progressively increased task difficulty by making stimuli smaller and lower in contrast (see Methods). Increasing task difficulty presented greater opportunity to examine trial-by-trial fluctuations of perception. Mice typically learned this task in 2-3 weeks, and performed hundreds of interleaved trials of monocular and binocular detection per day.

**Figure 1.**
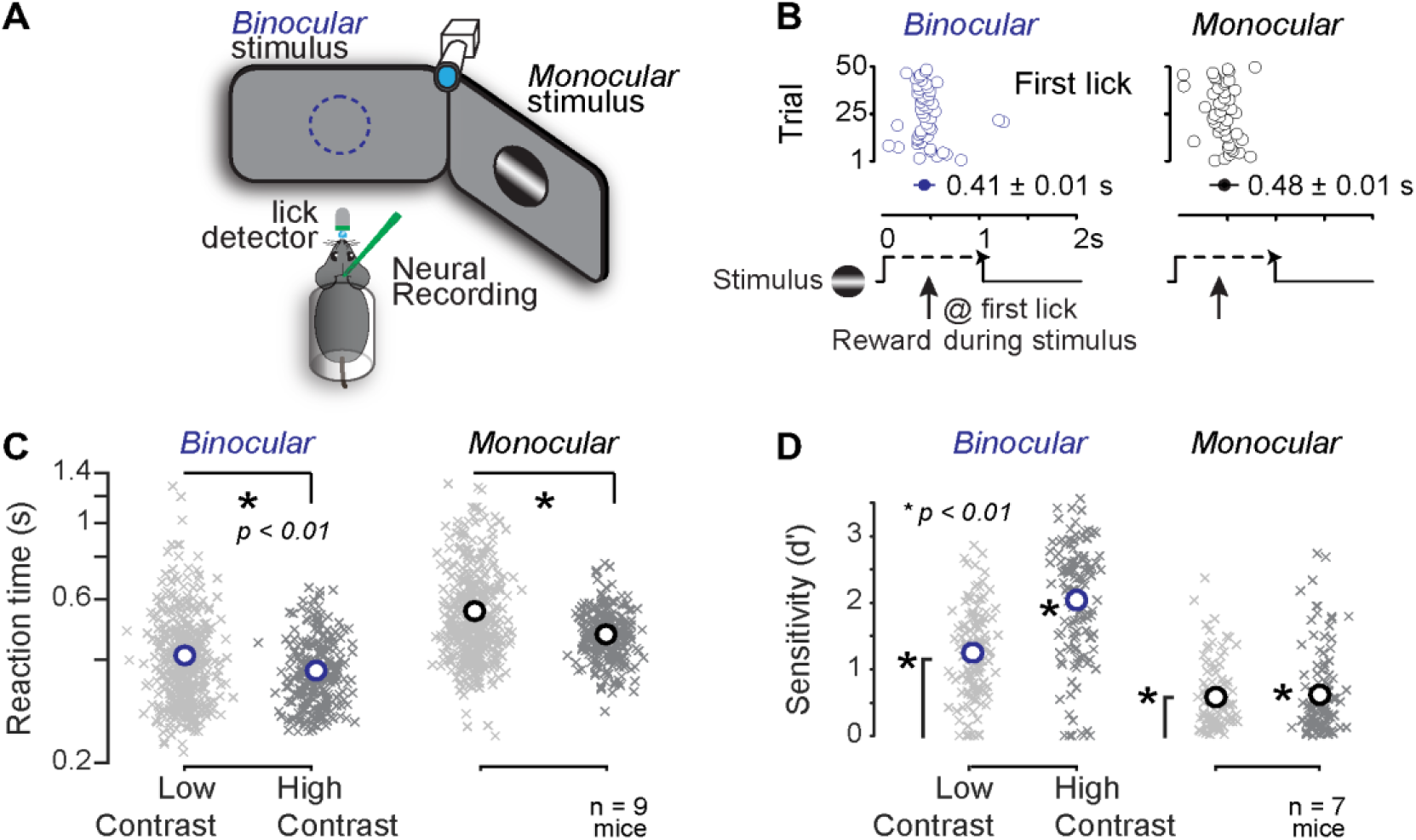
Visual detection latency and sensitivity depend upon spatial location. **A.** Head-fixed mice faced two monitors placed in the binocular (blue) and monocular (black) visual fields. Mice reported detection of visual stimuli (gratings) by licking for water. A photosensor recorded licks, a camera monitored the right eye, and silicon probes recorded activity in the left hemisphere. **B.** Example session of visual detection, where stimuli appeared in one location for 50 consecutive trials, then switched to the other location for 50 trials. Mice obtained reward upon first lick (open circles) during the stimulus presentation window (1 s duration). Reaction times were faster for binocular (blue) versus monocular detection (black). **C.** Reaction times decreased with increased stimulus contrast. Crosses indicate mean reaction time per block of 10-50 trials, circles indicate population mean reaction time (n = 9 mice, 752 blocks, >30k trials; last quartile of total training sessions). Binocular (blue): low contrast, 0.41 ± 0.04 s; high contrast: 0.37 ± 0.03; mean ± SEM; p < 0.01 rank sum test. Monocular (black): low contrast, 0.56 ± 0.06 s; high contrast, 0.47 ± 0.03; mean ± SEM; p < 0.01 rank sum test. Binocular low and high contrasts (mean): 45% and 67%; Monocular low and high contrasts (mean): 72% and 85%. **D.** Detection sensitivity (d’) improved with higher stimulus contrast. Crosses indicate d’ per daily session (n = 7 mice, 475 sessions, >115k trials from last quartile of training days). Circles indicate population mean. Binocular (blue): low contrast, 1.25 ± 0.06; high contrast: 2.04 ± 0.08; mean ± SEM; Monocular (black): low contrast, 0.59 ± 0.05; high contrast, 0.62 ± 0.06; mean ± SEM. All population means significantly greater than chance level (p < 0.01, sign test). Binocular sensitivity varied significantly with contrast (p < 0.01, rank sum test). Monocular sensitivity not different with contrast (p = 0.4). Binocular contrast means: 32% and 60%; Monocular contrast means: 83% and 98%.

Mice performed this detection task using vision. Stimulus location and contrast significantly affected reaction times and detection sensitivity. Mice detected binocular stimuli significantly more rapidly than monocular stimuli (Fig. 1C, left versus right panels). Moreover, within a given spatial location, higher contrast stimuli elicited significantly faster reaction times (Fig. 1C). Detection sensitivity (d’, see Methods) was significantly greater than chance level in both monocular and binocular visual fields, with the latter exhibiting greatest sensitivity (Fig. 1D). Lower visual contrast decreased detection sensitivity in both locations, significantly in the binocular visual field (Fig. 1D). Sensitivity to stimulus location and contrast did not depend upon grating orientation (not shown). Taken together, these results show that two major aspects of vision—spatial location and contrast — significantly influence visual detection behavior in mice.

Activity in primary visual cortex (V1) was necessary for stimulus detection. Pharmacological or optogenetic inactivation of monocular V1 abolished monocular detection, while interleaved trials of binocular detection were not significantly impaired during the same experiments (Fig. S1, see Methods). Inactivation of adjacent non-visual cortex caused no behavioral impairment. These results indicate that localized activity in V1 supports stimulus detection in retinotopically matched regions of visual space.

### Cortical states selectively modulate L4 LFP during visual spatial detection

We performed acute recordings of laminar population activity in V1 during visual spatial detection. We recorded in monocular V1 for two reasons. First, monocular detection trials exhibited greatest task difficulty; second, monocular stimuli activate V1 unilaterally, restricting the early stimulus-evoked activity to one hemisphere. We recorded from task relevant V1 neurons by measuring the spatial receptive field (RF) at each recording site, and ensuring that these overlapped the average location of the monocular stimuli during spatial detection (Fig. S2A).

LFP was starkly different during successful versus failed detection. Detection failures (Misses) were often accompanied by synchronized, low frequency (3 – 7 Hz) LFP oscillations during the stimulus (Fig. 2A, grey). By functionally identifying cortical layers (Niell and Stryker, 2008; Pluta et al., 2015), we found that these 3 – 7 Hz oscillations were strongest in Layer 4 (L4) and L5/6 (Fig. S2B-F). Moreover, 3 – 7 Hz residual LFP power was selectively and significantly elevated only during failed detection (Miss) trials (Fig. 2B-D).

**Figure 2.**
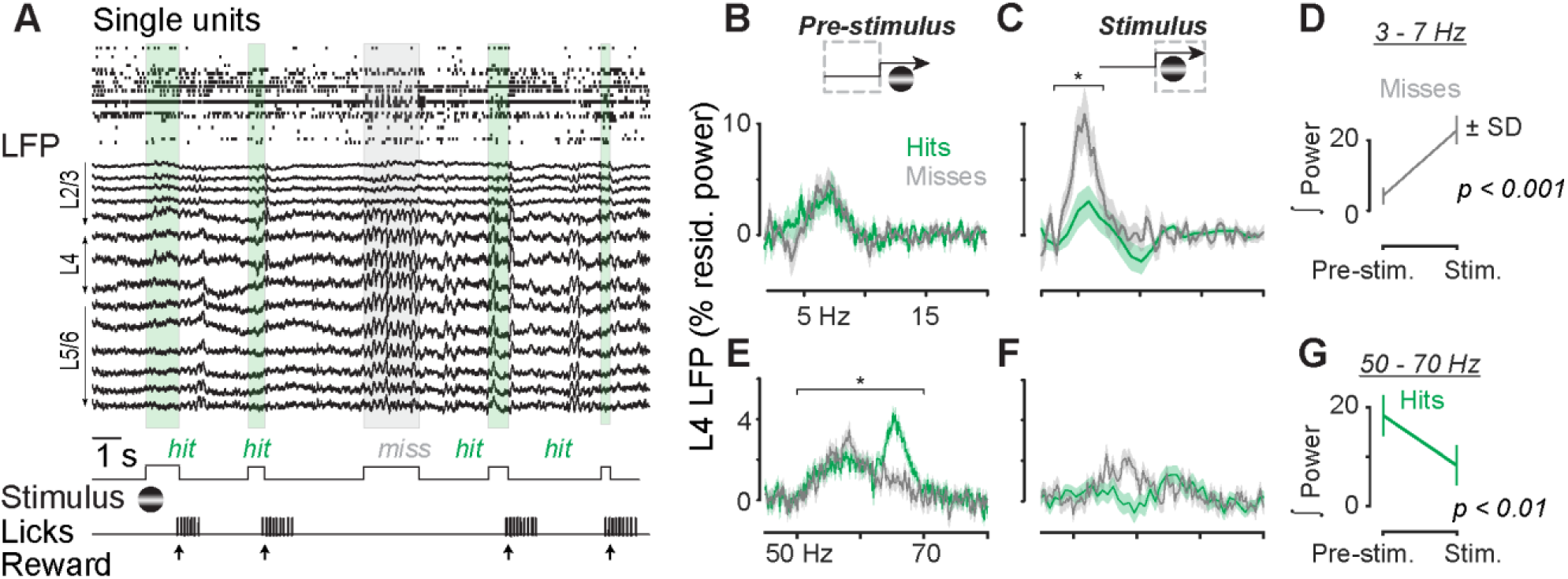
Cortical state signatures in layer 4 LFP during visual spatial detection. **A.** Single units and local field potential (LFP) recorded simultaneously with a multi-site probe in monocular visual cortex (V1) during visual detection. Layers 2/3 (L2/3), L4, and L5/6 estimated by current source density analysis. Detection failure (miss, grey box) occurred in the midst of successful detection trials (hits, green boxes). **B-C.** L4 LFP residual power in 2 – 20 Hz range on Hit (green) and Miss (grey) trials, measured for the pre-stimulus and stimulus periods. Traces indicate mean ± SEM, n = 15 sessions in 11 mice. Residual power integrated from 3 – 7 Hz significantly greater on Miss versus Hit trials during stimulus (paired signed rank test, p < 0.01). No significant difference between Hits and Misses pre-stimulus (p > 0.4). **D.** Significantly elevated 3 – 7 Hz integrated power on Miss trials during stimulus (paired signed rank test, p < 0.01). **E-F.** L4 LFP residual narrowband gamma (50 – 70 Hz) power on Hit (green) and Miss (grey) trials, measured for the pre-stimulus and stimulus periods. Residual power integrated from 50 – 70 Hz significantly greater on Hit versus Miss trials before stimulus (paired signed rank test, p < 0.01). **G.** Significantly reduced 50 – 70 Hz integrated power on Hit trials during stimulus (paired signed rank test, p < 0.01).

Successful detection was preceded by elevated narrowband gamma (50-70 Hz) LFP in L4. Narrowband gamma residual power was strongest in L4 (Fig. S2E), and in the absence of visual contrast. Remarkably, L4 narrowband gamma power varied with behavioral outcome: it was significantly elevated and then suppressed by the onset of visual contrast selectively on Hit trials (Fig. 2E-G).

### Cortical states decorrelate neurons and suppress firing before stimulus detection

We next examined laminar activity of single neurons comprising two distinct classes: broad waveform regular spiking (RS) putative excitatory neurons, and narrow waveform fast-spiking (FS) putative inhibitory neurons (Fig. 3A-B). Spike widths of FS neurons matched those of parvalbumin (PV) interneurons directly activated by channelrhodopsin (Fig. S3A-C). This suggests that FS neurons in our experiments are PV interneurons.

**Figure 3.**
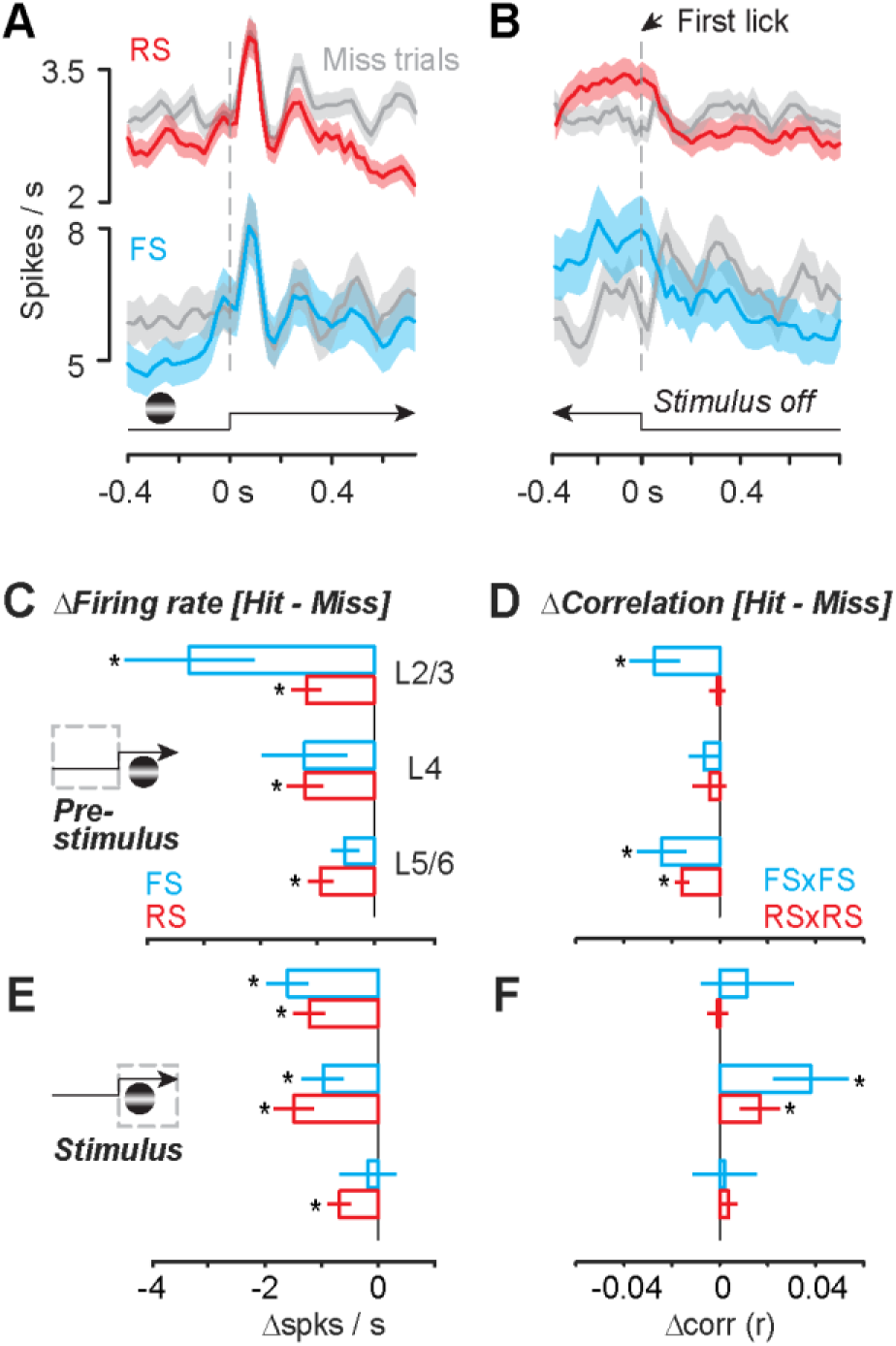
Cortical states decorrelate and suppress firing before stimulus detection. **A.** Regular Spiking (RS, n = 172) and Fast Spiking (FS, n = 52) neurons in monocular V1 during monocular detection (Hit trials, colored; Miss trials, grey). Mean ± SEM, aligned to stimulus onset. Spikes binned at 25 ms, smoothed ± 1 bin. Same experiments as Fig. 2. **B.** Same as in A, aligned either to first lick (Hit trials) or stimulus offset (Miss trials). **C.** RS cells fired significantly less in all layers before Hit trials (L2/3: −1.2 ± 0.3 spikes / s; L4: −1.2 ± 0.4; L5/6: −1.0 ± 0.2; mean ± SEM; cell-by-cell paired signed rank p < 0.01 for all; Fig. S3). FS cells fired significantly less in L2/3 before Hit trials (-3.3 ± 1.1 spikes / s), but not in L4 (−1.2 ± 0.8) or L5/6 (−0.5 ± 0.2; (p = 0.1 for both). **D.** FS pairs significantly decorrelated in L2/3 (−0.03 ± 0.01) and Layer 5/6 (−0.02 ± 0.01) before Hit trials (cell-by-cell paired signed rank p < 0.01; Fig. S7) but not in L4 (−0.007 ± 0.007). RS neuron pairs in L5/6 significantly decorrelated before Hit trials (−0.02 ± 0.003). No significant differences in RS correlations in L2/3 (−0.001 ± 0.004) or L4 (−0.004 ± 0.007). **E-F.** Same comparisons as C-D, during stimulus. RS cells fired significantly less in all layers on Hit trials (L2/3: −1.2 ± 0.3 spikes / s; L4: −1.5 ± 0.4; L5/6: −0.7 ± 0.2; p < 0.01 for all; Fig. S5). FS cells fired significantly less during Hit trials in L2/3 (−1.6 ± 0.4) and L4 (−1.0 ± 0.4) but not L5/6 (−0.2 ± 0.5). FS and RS neuron pairs significantly correlated in L4 (FS: 0.04 ± 0.02, RS: 0.02 ± 0.01; p < 0.01 for both), but not L2/3 (RS: 0.001 ± 0.004; FS: 0.01 ± 0.02) or L5/6 (RS: 0.004 ± 0.004; FS: 0.002 ± 0.01).

RS and FS neurons displayed two distinct activations during successful detection. The initial visual response peaked and terminated rapidly, hundreds of milliseconds before the average reaction time (Hit trials, Fig. 3A). By aligning to reaction times on Hit trials, a second, rapidly rising late phase response emerged prior to the first lick (Fig. 3B); this was not present on Miss trials aligned to stimulus offset (Fig. 3B, grey). Late phase activity on Hit trials was not a movement artefact: firing terminated abruptly upon reward delivery, even though mice continued to lick vigorously during reward consumption.

Lower firing rates and reduced correlations (decorrelation) preceded successful detection. On a cell-by-cell basis, pre-stimulus RS firing rates were significantly lower on Hit versus Miss trials, in all layers (Fig. 3C; Fig. S3D-F). L2/3 FS neurons also fired significantly less before stimuli on Hit trials (Fig. 3C). Accordingly, FS neuron pairs in L2/3 were significantly less correlated before successful detection trials (Fig. 2D), as were FS pairs and RS pairs in L5/6 (Fig. 2D). The effects of decorrelation and suppressed firing prior to successful detection were not simply driven by higher arousal: pupil dilates with arousal, but it was significantly smaller before successful detection (Fig. S4).

Surprisingly, undetected stimuli evoked the highest firing rates. On a cell-by-cell basis, undetected stimuli evoked significantly greater RS firing in all layers, and significantly greater FS firing in L4 and L2/3 (Fig. 3E; Fig. S5). LFP responses were also significantly greater for undetected stimuli (not shown). However, stimuli that were successfully detected increased RS and FS correlations selectively in L4 (Fig. 3F; Fig. S5), eliminating the decorrelated cortical state present before stimulus onset (cf. Fig. 3E).

### RS neurons and L4 LFP oscillations predict single-trial behavior most accurately

Which aspects of cortical states best predict visual spatial detection? We observed robust signatures of cortical states across network, laminar, and cellular levels, even when averaging across multiple behavioral sessions and subjects. We thus quantified the accuracy of predicting single trial behavior from cortical state signatures at both network (LFP) and cellular (RS / FS neuron) levels.

Prior to stimulus onset, RS neurons predicted single-trial behavior most accurately. RS firing rates were significantly more predictive than pairwise RS correlations (Fig. 4A; 68 ± 4% versus 62 ± 4% correctly predicted single trials; mean ± SD; p< 0.01; cross-validated linear classifier, see Methods). Both of these factors predicted better than pre-stimulus FS neuron activity, or network level LFP power. Prestimulus RS activity was also significantly more predictive than arousal level measured by pupil area (Fig. S4D; 60 ± 3%).

**Figure 4.**
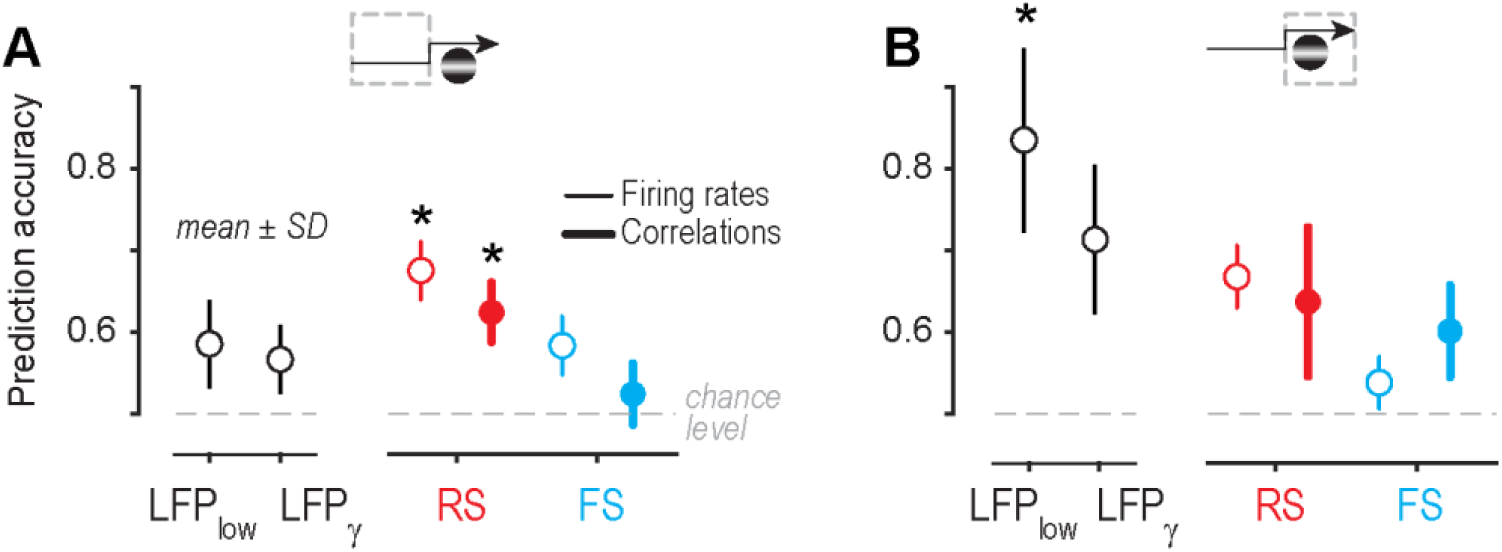
RS activity and L4 LFP oscillations predict single-trial behavior. **A.** Pre-stimulus RS firing rates predicted single trial behavior (68 ± 4% accuracy; mean ± SD) significantly better than RS correlations (62 ± 4%) FS firing rates (58 ± 4%) or FS correlations (52 ± 4%). Both RS rates and correlations were more predictive than network level factors (L4 LFP low frequency residual power: 59 ± 5%; L4 LFP narrowband gamma residual power), and pupil area (Fig. S4). Non parametric ANOVA with Tukey correction for multiple comparisons. All factors predicted greater than chance level (dashed line; p < 0.01, sign test for all). **B.** During the stimulus, L4 LFP low frequency residual power predicted single trials (84 ± 11%) better than all other factors. L4 LFP narrowband gamma predicted better (71 ± 10%) than the remaining factors. No significant difference between pre-stimulus versus stimulus-evoked RS rates (67 ± 4%) or correlations (64 ± 10%). Stimulus evoked FS correlations (60 ± 4%) predicted significantly better than pre-stimulus FS correlations (p < 0.01, rank sum test).

Upon stimulus onset, L4 LFP oscillations provided superior predictions of single-trial behavior. Stimulus evoked 3 – 7 Hz L4 LFP power predicted trial outcome with 84 ± 11% accuracy, better than any other stimulus-driven factor (Fig. 4B). In parallel, stimulus-evoked suppression of L4 narrowband gamma LFP power predicted 71 ± 10% of single trials, significantly better than the remaining factors. Remarkably, stimulus onset did not significantly improve predictions from RS firing rates and correlations (67 ± 4% and 64 ± 10%) compared to pre-stimulus predictions. Stimulus onset rendered FS neuron correlations significantly more predictive than pre-stimulus correlations (60 ± 4% vs 52 ± 4%, p < 0.01), but these remained less predictive than any form of RS activity.

## Discussion

Here we revealed that cortical states play a central role for visual spatial perception in mice. Cortical states — defined by decorrelation, suppressed firing, and elevated gamma power— accurately predicted single trials of visual behavior, even before stimulus onset. Our findings in mice recapitulate fundamental signatures of cortical states known to enhance visual perception in higher mammals.

Effects of cortical state on mouse visual perception are different from other sensory modalities. Cortical states with large prestimulus correlations and high firing rates were detrimental for visual detection. In stark contrast, somatosensory detection performance resists large fluctuations in cortical state, and does not depend upon the magnitude or frequency content of the early sensory response (Sachidhanandam et al., 2013). Our findings were not simply explained by arousal: pupil was largest, and firing rates highest, during failures of detection. Differences may arise from circuit organization across cortical areas, or from behavioral context. In our visual spatial task, monocular detection was more difficult than binocular detection, and this may have accentuated the effects of cortical state (Chen et al., 2008; McGinley et al., 2015; Spitzer et al., 1988). Our study isolated the effects of cortical state on perception in stationary conditions, minimizing complicated interactions between arousal, visual motion, and locomotion (Niell and Stryker, 2010; Poort et al., 2015; Saleem et al., 2013; Vaiceliunaite et al., 2013; Vinck et al., 2015). Understanding the effects of cortical states on perception across modalities, during a variety of behavioral contexts, remains an important topic for future study.

Selective coordination of population activity by cortical states supported visual perception in mice. First, elevated narrowband gamma power was a major factor predicting stimulus detection, similar to the role of gamma in visual detection in primates (Fries, 2015; Lima et al., 2011; Womelsdorf et al., 2006). Second, pre-stimulus decorrelation—most prominent in FS neurons— preceded successful stimulus detection, as in primates (Cohen and Maunsell, 2009; Mitchell et al., 2009). However, decorrelation and FS neuron activity did not predict behavior better than RS firing rates, at odds with findings in primates (Mitchell et al., 2007; Snyder et al., 2016). This may be due to differences in RS and FS neuron identification in higher mammals (Constantinople et al., 2009; Soares et al., 2017; Vigneswaran et al., 2011). In mice, >90% of FS neurons are PV inhibitory neurons, and >90% of RS neurons are excitatory neurons (Lee et al., 2010; Pfeffer et al., 2013; Rudy et al., 2011), enabling clearer interpretation of roles of cell types. Our findings have two important differences from primate studies. First, narrowband gamma in mouse V1 has origins and mechanisms distinct from broadband gamma (Saleem et al., 2017). Second, our effects on gamma and correlations were not directly elicited by visual attentional cues, as in primate studies.

Cortical states enabling perception exerted strong effects in the input layer of mouse V1. In L4, selective modulation of narrowband gamma and suppression of 3 – 7 Hz oscillations predicted 70-80% of single behavioral trials. A recent study of L2/3 neurons described similar low frequency oscillations, but these were less selective for behavioral outcomes (Einstein et al., 2017). Although these oscillations invaded all cortical layers, we found that they are prominent and strongly predictive of behavior in L4. In parallel, pre-stimulus narrowband gamma was strongest in L4, and the degree of its suppression strongly predicted behavior. Our study shows that cortical states exert widespread network and cellular effects in mouse V1, but these appear most predictive of visual behavior at the very first stage of visual cortical processing.

## Methods

### Contact for Reagent and Resource Sharing

All requests for resources should be directed to and will be fulfilled by Bilal Haider.

### Experimental model and subjects

All procedures were approved by the Institutional Animal Care and Use Committee at the Georgia Institute of Technology and were in agreement with guidelines established by the National Institutes of Health and the Animals (Scientific Procedures) Act 1986 (UK).

#### Surgery

Male C57BL6J mice (4-6 weeks old; reverse light cycle housing; bred in house; RRID: IMSR_JAX:000664) were chronically implanted with a custom-built stainless steel headplate with recording chamber (3-4 mm inner diameter) under isoflurane anesthesia (5% induction, 0.5% - 2% maintenance). The headplate was affixed to the skull using a thin layer of veterinary adhesive (VetBond) then securely bonded to the cranium (Metabond). The recording chamber was sealed with elastomer (KwikCast). Following implantation, mice were allowed to fully recover for 3 days. After recovery, animals were handled and acclimatized for 3-4 days to the head fixation apparatus, and then placed under a restricted water schedule for behavioral training. In some experiments, we performed similar procedures in male offspring of PV-cre (RRID: IMSR_JAX:017320) crossed with Ai-32-ChR2 (RRID: IMSR_JAX:024109) mice to optogenetically activate PV inhibitory neurons during behavior (Fig. S1).

#### Water restriction

Mice learned to perform visual detection for water rewards. Mice received a minimum daily amount of water (40 ml/kg/day; 1 ml/day for a typical 25 g mouse). Reference weights were computed according the previous methods (Burgess et al., 2017). Well-trained mice typically received all of their minimum daily water (and often more) exclusively during the task. Early in training, naïve mice often received less than their minimum amount of hydration in the task, so they received precisely measured supplemental hydration (hydrogel) to reach daily requirements.

### Behavior

#### Training

Mice learned the contingency between stimulus appearance and reward availability through passive instrumental conditioning. In early sessions, reward was automatically delivered after a fixed delay (0.6 −0.7 s) relative to stimulus onset. With training, the latency to first lick aligned to stimulus onset occurred before reward delivery, indicating that the mouse was reacting to the visual stimulus with anticipatory licks. The mouse then transitioned to active visual detection sessions, in which reward delivery occurred only if the mouse licked during the stimulus window (typically 1.5 - 2 seconds early in training, 1-1.5 s late in training; stimulus disappears upon first correct lick in stimulus window). Detection performance was quantified using the psychophysical sensitivity metric d-prime (d’; (Green and Swets, 1974). Hit rates were calculated from correct detection trials (licks during stimulus window) while false alarm rates were calculated from trials with blank targets (0% contrast; 20% of trials). When d’ was above chance levels for 2 consecutive days, stimulus contrast range and/or size was decreased to maintain difficulty. Once animals exhibited performance above chance for both binocular and monocular stimuli of high and low contrast, we performed acute neural recordings.

#### Stimulus shaping

Mice detected static horizontal Gabor gratings, presented in either the binocular or monocular visual field on linearized LCD monitors (60 or 80 Hz refresh rate). Grating phase was randomized every trial, while spatial frequency (range 0.05 - 0.1 cycles/deg) and stimulus size (σ range 10°-20°) remained fixed across blocks of trials. Binocular contrasts ranged from 2% - 75% (38 ± 26% during recording sessions, mean ± SD) monocular contrasts ranged from 50%-90% (74 ± 21%). During recording sessions, in addition to the detected stimuli, task-irrelevant stimuli were presented to facilitate receptive field mapping (bars 9 degrees wide, 2.5-5% contrast, 0.1 ms duration, 0.3 s interval, randomized location). These faint and brief bars did not affect behavior, and are not analyzed here.

#### Cortical inactivation

A glass micropipette (∼10 μm tip) was filled with 5 μL of 5 μg/μL muscimol (Sigma) dissolved in artificial cerebrospinal fluid. The pipette was lowered in monocular V1 to a depth of 700 μm (L5), and positive pressure of ∼17mBar ejected 0.1 μL over 6-10s. The pipette was slowly withdrawn to 300 μm, and the procedure was repeated in L2/3, as in prior studies (Komiyama et al., 2010). In optogenetic inactivation experiments with PV-cre x Ai32(ChR2) mice (Fig. S1), the skull was thinned over monocular V1 and a fiber-coupled LED (473 nm) delivered pulses of light to inactivate V1 on 25% of detection trials (1 s during visual stimulus, starting 0.1 s before stimulus and ramping to 5.8 mW measured at the cranium).

### Recordings

#### Surgical preparation

On the day of recording, mice were anesthetized with isoflurane and a small (∼100-500 μm) craniotomy was opened over the monocular portion of V1 (0.5 mm anterior to lambda, 2-2.5 mm lateral to central suture). Mice recovered for at least 3 hours prior to recording during behavior.

#### Electrophysiology

Recordings were done with multi-site silicon probes (NeuroNexus) consisting of either a single 32-channel shank, or two 16-channel shanks. Electrodes were advanced ∼ 1000 μm below the cortical surface. Data were collected for the duration of a behavioral session (typically 100-200 trials), after which a task-irrelevant stimulus was presented to facilitate receptive field mapping of the recording site (100% contrast vertical flashed bars, 9 degrees in width, duration 0.1 s, inter-stimulus interval 0.3 s, placed in random locations tiling 144 degrees of the visual field). The craniotomy was kept sterile and covered with elastomer in between consecutive recording days (typically 2-4 from the same site).

#### Eye Tracking

In a subset of mice, we simultaneously recorded the pupil (6 mice across 90 sessions, 5722 trials). A high speed camera (Imaging source DMK 21Bu04.H) with a zoom lens (Navitar 7000) and infrared filter (Mightex, 092/52x0.75) was placed approximately 22 cm from the animal’s right eye under near-infrared LED illumination (Mightex, SLS-02008-A). Video files were acquired and processed using the Image Acquisition Toolbox in MATLAB. 1 mm corresponded to ∼74 pixels on each frame.

### Analysis

The total neural data set consisted of 15 recording sessions from 11 mice, 1175 hit trials, 1031 miss trials, n = 224 units (52 FS cells,172 RS cells). All details of statistical comparisons are contained in the figure legends.

#### Spike sorting

Raw electrical signals were amplified and digitized (Blackrock Microsystems) then exported for post processing. Extracellular spikes were isolated using the KlustaViewa Suite (Rossant et al.,2016). Briefly, automated clustering was followed by three manual steps. First, obvious noise artifacts were eliminated. Second, poorly isolated waveforms were classified as multiunit activity. Waveforms of the remaining clusters were carefully curated in PCA space in parallel with unit auto- and cross-correlation histograms to define well-isolated single units. A small number of units (n = 7) with low signal to noise ratio (SNR) < 3 were excluded because of unreliable measurement of spike widths. No additional criteria were used to include or exclude units from further analysis. The average SNR of our units (n = 224) was 35.0, and the average recording yield was 17.2 ± 8.9 units per session (mean ± SD).

We classified Fast spiking (FS) and Regular Spiking (RS) units according to spike width. Histograms of population spike widths (measured peak to trough) were clearly bimodal (Fig. S3). Units with a peak-to-trough width less than 0.57 ms were classified as FS, and broader units classified as RS. This classification of FS neurons closely agrees with previous studies of mouse V1 (Niell and Stryker, 2010), where FS neurons consist nearly exclusively of parvalbumin (PV) positive inhibitory neurons (Pfeffer et al., 2013). We additionally verified the inhibitory identity of FS neurons in our awake recording conditions by expressing channelrhodopsin in parvalbumin (PV) interneurons, and measuring spikes of PV interneurons directly activated by light (Fig. S3).

#### Correlations

Spike trains were convolved with a 20ms Gaussian filter, and cross correlations were calculated between smoothed spike trains (MATLAB xcorr function with ‘coeff’ parameter for normalization). Pre-stimulus period spike trains were limited to 3 seconds (or less, if the inter-trial interval was smaller on that particular trial, range: 0.5 – 3 s). Stimulus period spike trains consisted of the 0.5 seconds preceding the stimulus onset until the reaction time for correct trials, or until stimulus offset (1 - 2 seconds) for failed detection trials.

#### Classifier

We trained a support vector machine (SVM) classifier to predict behavioral outcome from simultaneously recorded neural activity (1175 hit trials, 1031 miss trials). Training sets consisted of firing rates or correlations sub-sampled in a ratio and sum that matched the average recording session (5 FS cells and 15 RS cells). For layer specific correlations, we sub-sampled a total of 15 neuron pairs within specific layers per trial, again to match statistics of the average recording session. Training and testing sets consisted of 200 trials each (100 hit and 100 miss). Gaussian kernels were Bayes optimized (MATLAB ’bayesopt’ function), and classifier performance was determined by the fraction of correctly classified trials in the testing set (MATLAB ‘fitcsvm’ function). This entire procedure was repeated 50 times to compute the mean and standard deviation of classifier performance for the dataset.

In a similar way, we constructed a classifier with the time series of pupil diameter in the 3 seconds preceding the stimulus. Again, training and testing sets consisted of 100 randomly selected hit and miss trials, and performance was determined by fraction of correctly classified trials in the testing set.

Trial classification with LFP gamma power used the residual power spectrum of the LFP between 50-70Hz, and low frequency LFP power 5 – 10 Hz, with the same training and testing parameters described above.

#### Pupil analysis

We first sub-selected a region of interest that captured changes in pupil for all frames in the file (Fig. S4A). Frames were then smoothed using a 2D Gaussian filter. We then selected intensities to separate the pixels within and outside of the pupil. A contour was drawn according to the identified intensities, and a least squares error 2D ellipse was fit to the contours. Pupil area was calculated as the percent deviation from the mean 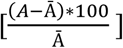, where A is the area in pixels and Ā is the average area across all frames. If an ellipse could not be fit, a NaN value was inserted for that frame. For analysis of pupil position, we looked at changes in the azimuthal coordinate of the fitted ellipse across frames, since this was the axis in which stimulus position varied.

#### LFP analysis

Local field potentials were obtained by bandpass filtering raw neural signals from 0.01 to 200 Hz. Laminar LFP responses were calculated by averaging across channels within an identified layer. We then averaged laminar LFP responses across behavioral sessions and animals (11 mice, 15 sessions, 2206 trials) to evaluate differences for hit and miss trials.

We analyzed the excess (residual) LFP power during Hit and Miss trials in low (2-20 Hz) and narrowband gamma (50-70 Hz) frequencies. As in our previous studies, we calculated residual power by fitting the entire power spectrum with a single exponential that excluded the band of interest. The difference (actual – fit) is divided by the fit to obtain fractional residual power. The power spectral density (FFT of the autocorrelation of the LFP) was calculated during the pre-stimulus and stimulus presentation epochs. Narrowband gamma (50-70 Hz) residual power was estimated by fitting the spectral density between 30-90 Hz, excluding 50-70 (Saleem et al., 2017). Residual low frequency power was estimated in the same manner, by fitting between 2-20 Hz excluding 4-12 Hz. Residual power was calculated per trial, then averaged across sessions and mice.

## Data availability

All data structures and code that generated each figure are available upon reasonable request.

## Author contributions

A.S., J.D.R., C.P.B., B.H. performed experiments, C.P.B. wrote software, A.S., J.D.R., B.H. performed analysis, B.H. and A.S. wrote the first draft with input from co-authors.

## Acknowledgements

We thank Aman Saleem for comments on an earlier version of this work, Tatsuo Sato, Michael Krumin, Charu Reddy, Alexander Zorn, and Hayley Arrowood for technical assistance, and Matteo Carandini for support in the initial phase of this study (Wellcome Trust 095668 and 095669). J.D.R. was funded by a Goizueta Foundation fellowship. B.H. was funded by the National Science Foundation (IRFP 0965110), the GT Neural Engineering center (1241384), and the Whitehall Foundation.

## SUPPLEMENTARY FIGURES

**Fig. S1.**
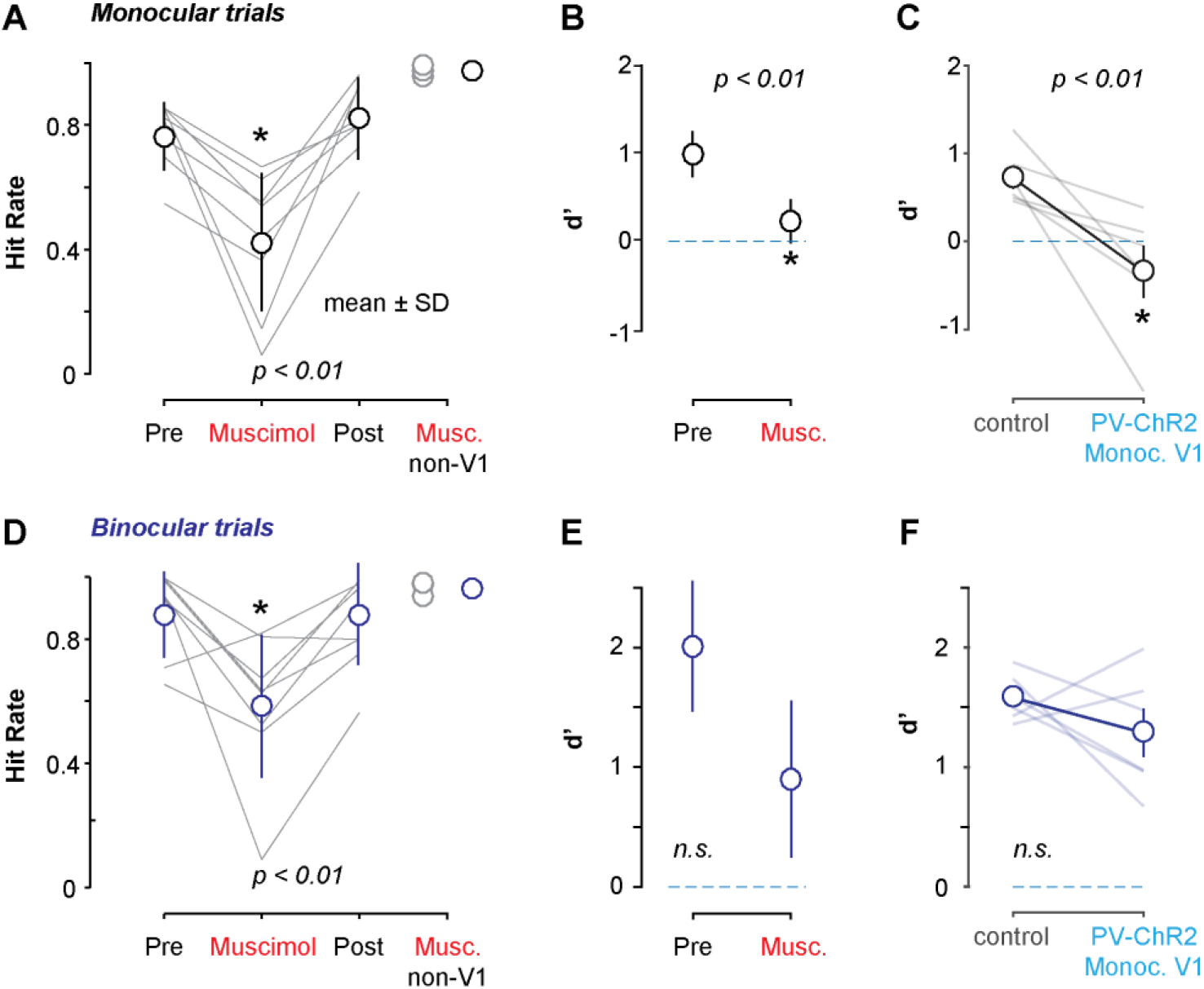
Inactivation of monocular V1 selectively impairs monocular visual detection (related to Fig.1). **A.** Intracortical injection of muscimol into monocular V1 significantly impaired monocular detection (Hit rates: 76% ± 11% versus 42 ± 22%; mean ± SD; n = 4 mice, 8 sessions, 2055 trials; p < 0.01 Wilcoxon signed-rank test). Hit rates recovered the next day (82 ± 11%). Inactivation of adjacent cortex (retrosplenial or parietal) did not affect hit rates, nor did injection of artificial cerebrospinal fluid in V1 (n = 1 mouse, 3 sessions, 1151 trials; not shown). Injection sites were mapped for spatial selectivity before inactivation (Fig. S3). See Methods for inactivation protocol. **B.** Inactivation significantly reduced monocular detection sensitivity (d’; 0.97 ± 0.3 vs. 0.21 ± 0.3; mean ± SD); p < 0.01 Wilcoxon signed-rank test). Sensitivity during inactivation is not significantly different from chancel level (p = 0.3, sign test). **C.** Optogenetic inhibition of monocular V1 (interleaved on 25% of trials) significantly reduced monocular d’ (0.72 ± 0.13 to −0.34 ± 0.3; mean ± SD; p < 0.05; Wilcoxon signed-rank test; n = 6 days and 1500 trials). Data from one PV-cre x A1-32-ChR2 mouse. **D-E.** Same sessions as **A-B**, muscimol inactivation significantly reduced binocular hit rates (88 ± 14% versus 59 ± 23%, p < 0.01, Wilcoxon signed-rank test), but did not significantly reduce sensitivity compared to control conditions (d’; 2.0 ± 0.5 vs. 0.9 ± 0.7; p = 0.1, Wilcoxon signed-rank test), and d’ remained significantly above chance levels (p < 0.01, sign test). **F.** Same sessions as **C,** optogenetic inhibition had no significant effect on binocular detection (d’: 1.58 ± 0.08 to 1.29 ± 0.2; p = 0.69).

**Fig. S2.**
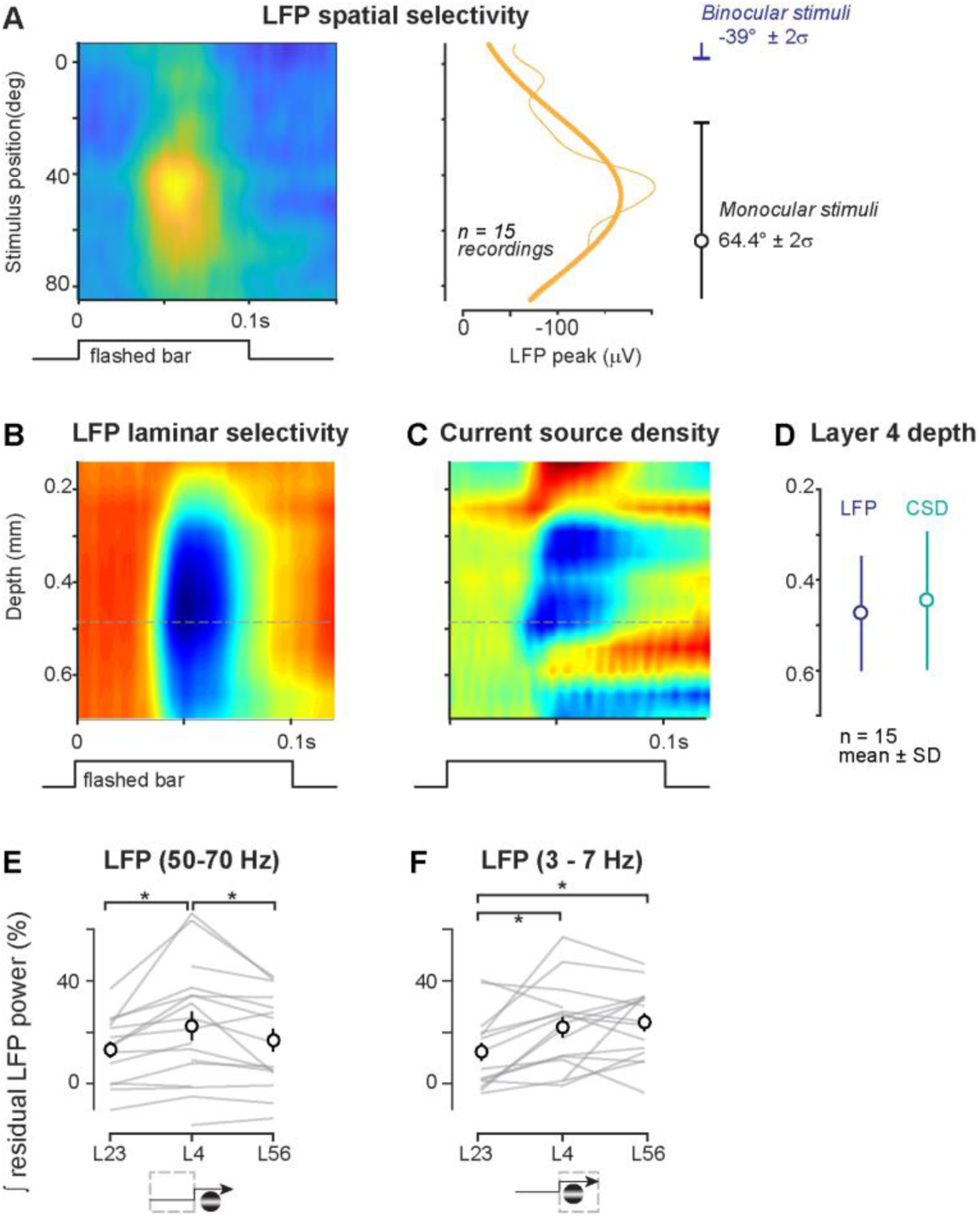
Spatial tuning and laminar analysis of LFP residual power (related to Fig. 2). **A.** Left, average space-time receptive field (RF) maps of local field potential (LFP) responses across the population (n = 15 recordings). Right, peak LFP responses (thin line) and Gaussian fit (thick line). LFP responses peak at 44 ± 52° (mean ± 2σ of fit), monocular stimulus presented at 64 ± 30° (mean ± 2σ of Gabor). Note that the Binocular stimulus was nearly 100° away. **B.** Example recording of average laminar LFP response to high contrast bars flashed at the center of the receptive field. Dashed line indicates depth of maximum LFP response. 32 sites spanning 775 microns. Data interpolated and smoothed for display, see Methods. **C.** Current source density (CSD) calculated for recording in A. Earliest and largest current sink corresponds to the site of maximum LFP response (dashed line, as in A). **D.** Estimated Layer 4 depth from LFP peak (466 ± 123 μm, blue) versus CSD sink (453 ± 154 μm, teal). Both depth estimates agree with prior studies (Lien and Scanziani, 2013; Pluta et al., 2015) **E.** Pre-stimulus narrowband gamma (50 – 70 Hz) residual power across layers. All electrode contacts within a layer were averaged within individual experiments (lines), then averaged across experiments (black, mean ± SEM, n = 17). A few recordings used electrodes that did not span all layers. Residual narrowband gamma power in L4 (0.22 ± 0.06) significantly greater than L2/3 (0.13 ± 0.03; p < 0.001) and L5/6 (0.17 ± 0.04; p < 0.01). L2/3 and L5/6 not significantly different (p = 0.09; one-tailed paired signed rank tests for all). **F.** Stimulus-evoked low frequency (3 – 7 Hz) residual power across layers. Same experiments as A. L4 residual power (0.22 ± 0.04) significantly greater than L2/3 (0.12 ± 0.04; p < 0.01). L4 and L5/6 (0.24 ± 0.04) not significantly different (p = 0.68). L5/6 significantly greater than L2/3 (p = 0.01; one-tailed paired signed rank tests for all).

**Figure S3.**
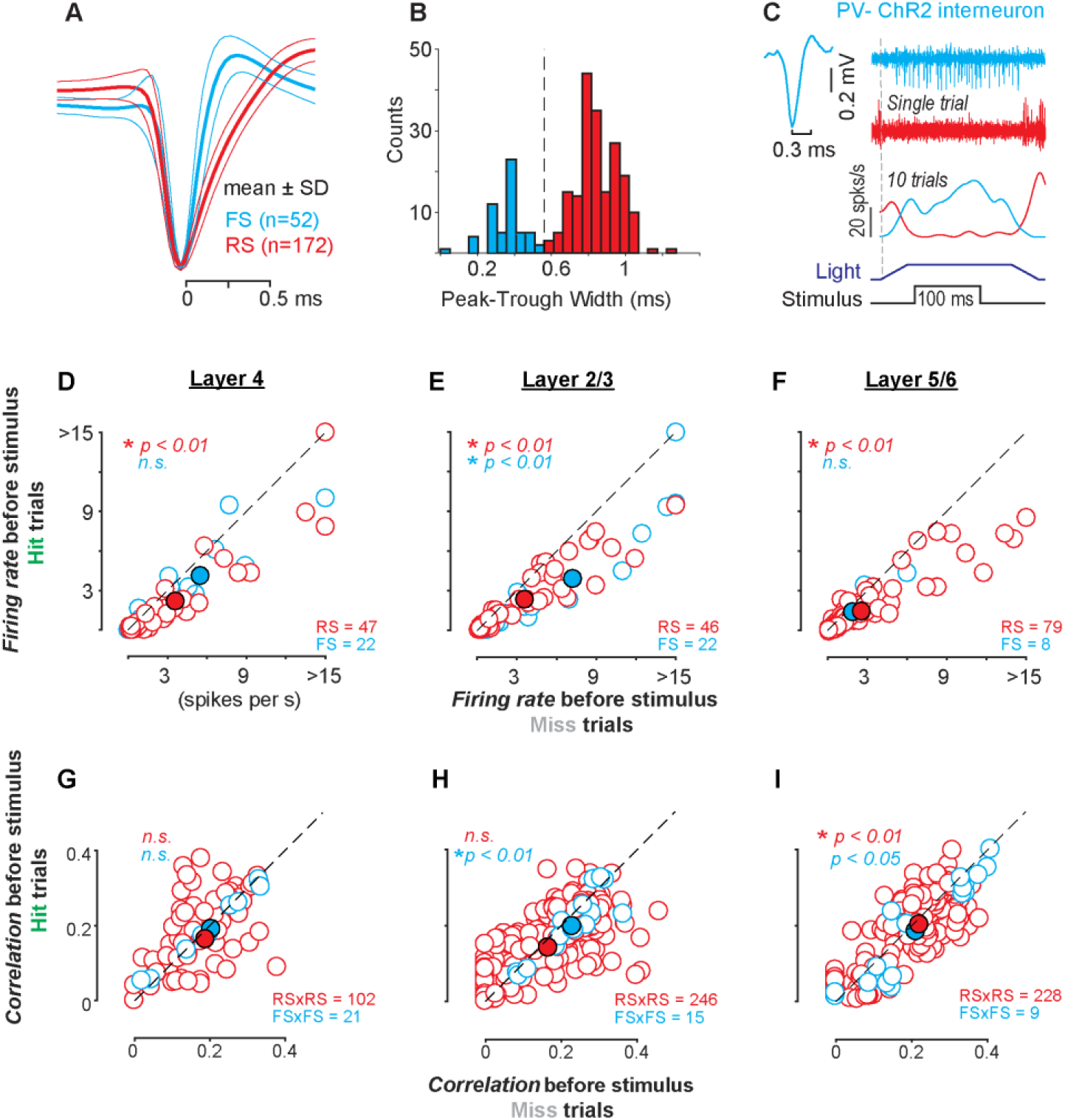
Pre-stimulus activity of fast spiking (FS) and regular spiking (RS) neurons during visual spatial detection (related to Fig. 3). **A-B.** Mean waveforms of RS (red, n = 172) and FS (cyan, n = 52) neurons (± SD), and histogram of all peak to trough spike widths. Individual waveforms normalized to minimum value before averaging. **C.** PV interneuron spikes directly activated by photostimulation of channelrhodopsin (bottom). PV spike width (0.3 ms) overlaps with FS neurons in B. **D-F.** Pre-stimulus firing rates before detection failure (miss trials, abscissa) versus success (hit trials, ordinate). Layers identified with current source density (see Fig. S2). In all layers, regular spiking (RS) units fired significantly less before successful detection (L4: 3.4 ± 0.9 vs. 2.2 ± 0.8 spikes / s; L2/3: 3.6 ± 0.6 vs. 2.4 ± 0.4; L5/6: 2.5 ± 0.4 vs. 1.6 ± 0.2; mean ± SEM; p < 0.01, Wilcoxon signed-rank test for all). FS neurons fired significantly less before successful detection in L2/3 (7.2 ± 2.2 vs. 4.0 ± 1.3 spikes / s; p < 0.01, Wilcoxon signed-rank test), but not in other layers (L4: 5.5 ± 1.6 vs. 4.3 ± 1.3; p = 0.33; L5/6: 1.9 ± 0.7 vs. 1.4 ± 0.6; p = 0.33). Spikes counted from stimulus onset to reaction time (hits), or until stimulus offset (misses). **G-I.** Noise correlations before stimulus onset in RS neuron pairs (RS x RS, red) and FS neuron pairs (FS x FS, cyan) within layers. RS pairs significantly reduced noise correlations (decorrelated) prior to successful detection in L5/6 (0.20 ± 0.006 vs. 0.19 ± 0.006; mean ± SEM; p < 0.01, Wilcoxon signed-rank test for all), but not in other layers (L4: 0.18 ± 0.01 vs. 0.18 ± 0.008; p = 0.1 L2/3: 0.16 ± 0.006 vs. 0.16 ± 0.005; p = 0.4). FS pairs significantly reduced noise correlations in L2/3 and L5/6 (L2/3: 0.23 ± 0.02 vs. 0.20 ± 0.02; p < 0.01; L5/6: 0.21 ± 0.03 vs. 0.19 ± 0.03; p < 0.05), but not L4 (L4: 0.20 ± 0.04 vs. 0.19 ± 0.03; p = 0.2).

**Figure S4.**
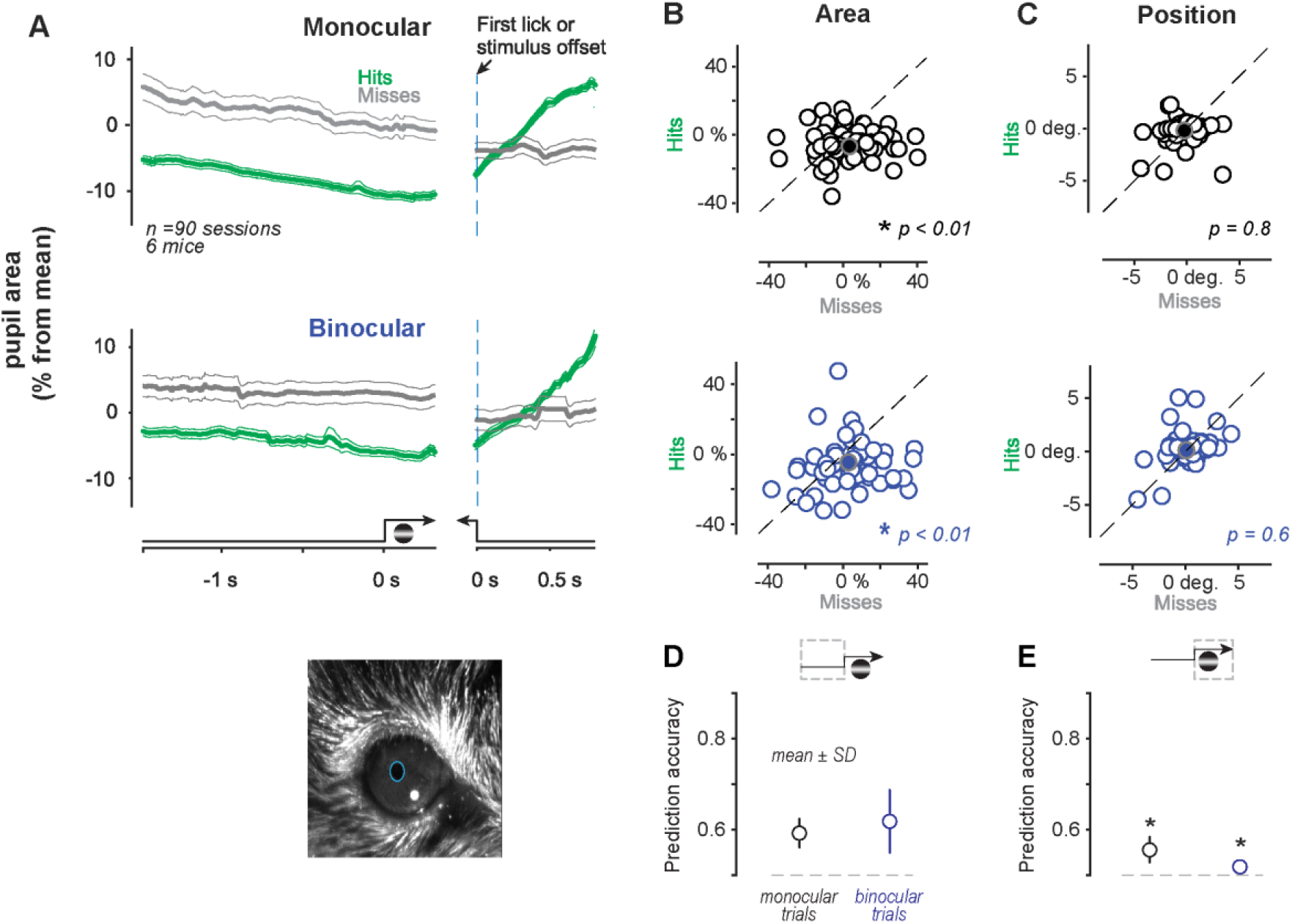
Pupil dynamics and single-trial predictions of visual spatial detection (related to Fig. 4) **A.** Pupil area (Δ% from mean of entire session) before successful detection (Hits, green) of monocular (top rows) and binocular stimuli (bottom rows). Pupil dilates rapidly after the first lick and reward on all Hit trials. Monocular grating stimulated the imaged eye. Same experiments with interleaved trials. Mean ± SEM, 90 sessions in 6 mice. **B.** Pupil area was significantly smaller before Hits in both monocular (top, 4 ± 13%; mean ± SD; p < 0.01) and binocular trials (bottom, 3 ± 16%; p < 0.01 for both, Wilcoxon signed-rank test). **C.** Pre-stimulus pupil position was not significantly different before Hits versus Misses in either monocular (0.5 ± 2.3° versus −0.5 ± 2.3°; mean ± SD; p = 0.8) or binocular trials (−0.1 ± 2.3°, −0.1 ± 2.6°; p = 0.6,Wilcoxon signed-rank test). **D.** Pre-stimulus pupil area predicted single trial detection of monocular (60 ± 3% accuracy; mean ± SD) and binocular (62 ± 7%) stimuli significantly better than chance (dashed line; p < 0.01, sign test for both). Pre-stimulus predictions were slightly but significantly better on binocular versus monocular trials (p = 0.02; rank sum test). **E.** Pupil area was significantly less predictive during the stimulus (monocular: 56 ± 3%; binocular: 52 ± 1%; p < 0.001 rank sum tests; pre-stimulus versus stimulus, within location). Moreover, during the stimulus, predictions were superior for monocular versus binocular detection (p < 0.001, rank sum test).

**Figure S5.**
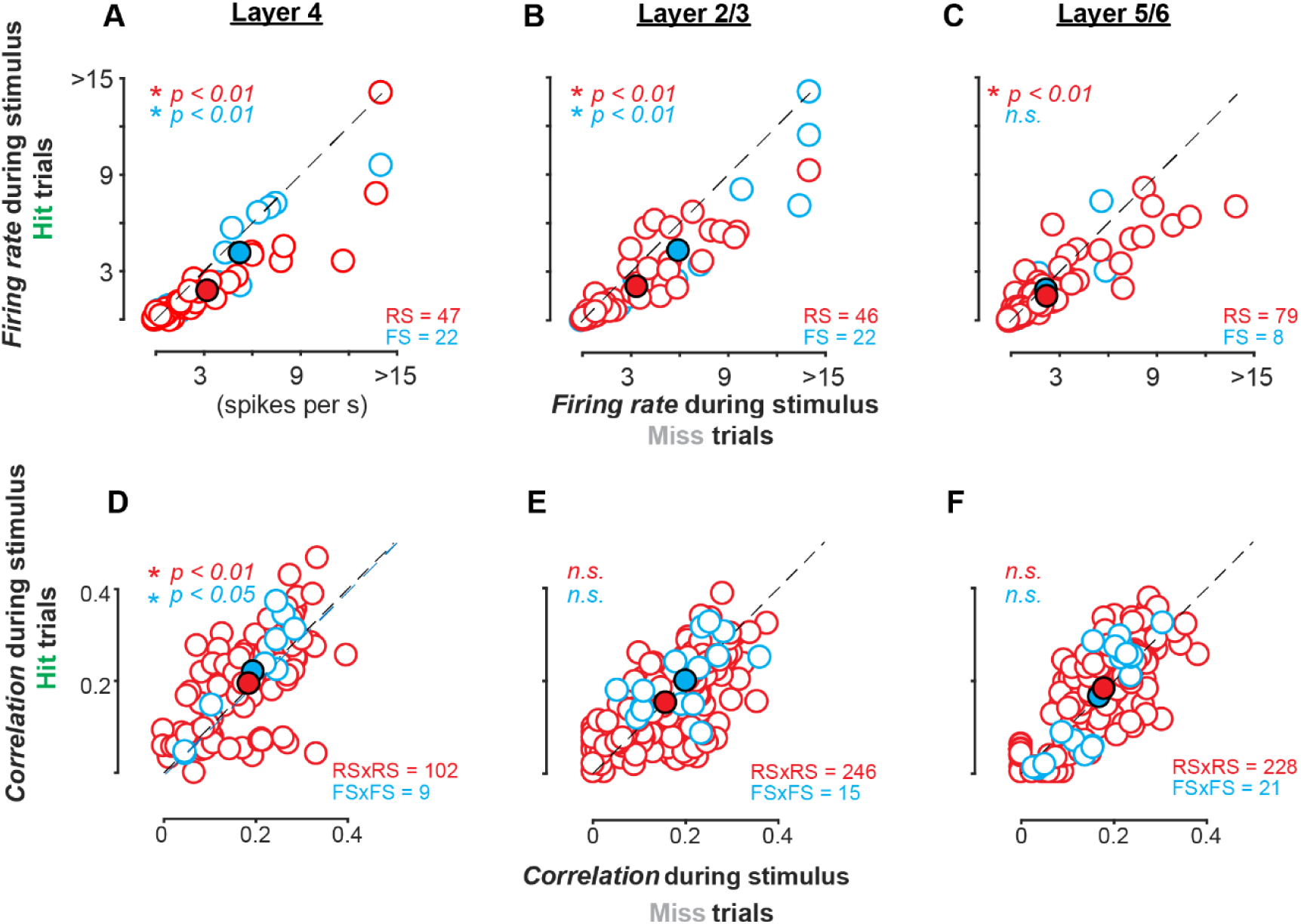
Stimulus-evoked activity of RS and FS neurons during visual spatial detection (related to Fig.3) A-C. Stimulus evoked firing rates for detection failure (Miss trials, abscissa) versus success (Hit trials, ordinate). Layers identified with current source density (see Fig. S4). In all layers, regular spiking (RS) units fired significantly less during Hits versus Misses (L4: 2.0 ± 0.6 versus 3.5 ± 0.9 spikes / s; L2/3: 2.4 ± 0.4 versus 3.6 ± 0.5; L5/6: 1.7 ± 0.2 versus 2.4 ± 0.4 spikes / s; mean ± SEM; p < 0.01, Wilcoxon signed-rank test for all). In L4 and L2/3, FS neurons fired significantly less during Hits versus Misses (L4: 4.6 ± 1.4 versus 5.5 ± 1.6 spikes / s; L2/3: 4.8 ± 1.8 versus 6.4 ± 1.9 spikes / s; p < 0.01, Wilcoxon signed-rank test; L5/6: 2.2 ± 0.9 versus 2.4 ± 0.9 spikes / s; p = 0.5). Spikes counted from stimulus onset to reaction time (Hits), or until stimulus offset (Misses). **D-F.** Noise correlations in L4 RS pairs (RS x RS, red) significantly increased during successful detection (L4: 0.2 ± 0.01 versus 0.19 ± 0.01; p < 0.01; L2/3: 0.15 ± 0.005 versus 0.15 ± 0.006; p = 0.1; L5/6: 0.18 ± 0.007 versus 0.17 ± 0.006; p = 0.07; mean ± SEM; Wilcoxon signed-rank test for all). Likewise, L4 FS pairs (FS x FS, cyan) significantly increased noise correlations during successful detection (L4: 0.22 ± 0.04 versus 0.19 ± 0.03; p < 0.05; L2/3: 0.20 ± 0.02 versus 0.19 ± 0.02; p = 0.2; L5/6: 0.17 ± 0.03 vs. 0.17 ± 0.02; p = 0.4).

## References

Burgess, C.P., Lak, A., Steinmetz, N.A., Zatka-Haas, P., Bai Reddy, C., Jacobs, E.A.K., Linden, J.F., Paton, J.J., Ranson, A., Schroder, S., et al. (2017). High-Yield Methods for Accurate Two-Alternative Visual Psychophysics in Head-Fixed Mice. Cell Rep 20, 2513–2524.

Chen, Y., Martinez-Conde, S., Macknik, S.L., Bereshpolova, Y., Swadlow, H.A., and Alonso, J.M. (2008). Task difficulty modulates the activity of specific neuronal populations in primary visual cortex. Nat Neurosci 11, 974–982.

Cohen, M.R., and Maunsell, J.H. (2009). Attention improves performance primarily by reducing interneuronal correlations. Nat Neurosci 12, 1594–1600.

Constantinople, C.M., Disney, A.A., Maffie, J., Rudy, B., and Hawken, M.J. (2009). Quantitative analysis of neurons with Kv3 potassium channel subunits, Kv3.1b and Kv3.2, in macaque primary visual cortex. J Comp Neurol 516, 291–311.

Einstein, M.C., Polack, P.O., Tran, D.T., and Golshani, P. (2017). Visually Evoked 3-5 Hz Membrane Potential Oscillations Reduce the Responsiveness of Visual Cortex Neurons in Awake Behaving Mice. J Neurosci 37, 5084–5098.

Engel, T.A., Steinmetz, N.A., Gieselmann, M.A., Thiele, A., Moore, T., and Boahen, K. (2016). Selective modulation of cortical state during spatial attention. Science 354, 1140–1144.

Fries, P. (2015). Rhythms for Cognition: Communication through Coherence. Neuron 88, 220–235.

Gilbert, C.D., and Li, W. (2013). Top-down influences on visual processing. Nat Rev Neurosci 14, 350–363.

Green, D.M., and Swets, J.A. (1974). Signal detection theory and psychophysics (Huntington, N.Y.,: R. E. Krieger Pub. Co.).

Haider, B., Krause, M.R., Duque, A., Yu, Y., Touryan, J., Mazer, J.A., and McCormick, D.A. (2010). Synaptic and network mechanisms of sparse and reliable visual cortical activity during nonclassical receptive field stimulation. Neuron 65, 107–121.

Haider, B., and McCormick, D.A. (2009). Rapid neocortical dynamics: cellular and network mechanisms. Neuron 62, 171–189.

Hansen, B.J., Chelaru, M.I., and Dragoi, V. (2012). Correlated variability in laminar cortical circuits. Neuron 76, 590–602.

Harris, K.D., and Thiele, A. (2011). Cortical state and attention. Nat Rev Neurosci 12, 509–523.

Kohn, A., Coen-Cagli, R., Kanitscheider, I., and Pouget, A. (2016). Correlations and Neuronal Population Information. Annu Rev Neurosci 39, 237–256.

Kohn, A., Zandvakili, A., and Smith, M.A. (2009). Correlations and brain states: from electrophysiology to functional imaging. Curr Opin Neurobiol 19, 434–438.

Komiyama, T., Sato, T.R., O’Connor, D.H., Zhang, Y.X., Huber, D., Hooks, B.M., Gabitto, M., and Svoboda, K. (2010). Learning-related fine-scale specificity imaged in motor cortex circuits of behaving mice. Nature 464, 1182–1186.

Lee, S., Hjerling-Leffler, J., Zagha, E., Fishell, G., and Rudy, B. (2010). The largest group of superficial neocortical GABAergic interneurons expresses ionotropic serotonin receptors. J Neurosci 30, 16796–16808.

Lima, B., Singer, W., and Neuenschwander, S. (2011). Gamma responses correlate with temporal expectation in monkey primary visual cortex. J Neurosci 31, 15919–15931.

McGinley, M.J., David, S.V., and McCormick, D.A. (2015). Cortical Membrane Potential Signature of Optimal States for Sensory Signal Detection. Neuron 87, 179–192.

Mitchell, J.F., Sundberg, K.A., and Reynolds, J.H. (2007). Differential attention-dependent response modulation across cell classes in macaque visual area V4. Neuron 55, 131–141.

Mitchell, J.F., Sundberg, K.A., and Reynolds, J.H. (2009). Spatial attention decorrelates intrinsic activity fluctuations in macaque area V4. Neuron 63, 879–888.

Nandy, A.S., Nassi, J.J., and Reynolds, J.H. (2017). Laminar Organization of Attentional Modulation in Macaque Visual Area V4. Neuron 93, 235–246.

Niell, C.M., and Stryker, M.P. (2008). Highly selective receptive fields in mouse visual cortex. J Neurosci 28, 7520–7536.

Niell, C.M., and Stryker, M.P. (2010). Modulation of visual responses by behavioral state in mouse visual cortex. Neuron 65, 472–479.

Nienborg, H., Cohen, M.R., and Cumming, B.G. (2012). Decision-related activity in sensory neurons: correlations among neurons and with behavior. Annu Rev Neurosci 35, 463–483.

Petersen, C.C., and Crochet, S. (2013). Synaptic computation and sensory processing in neocortical layer 2/3. Neuron 78, 28–48.

Pfeffer, C.K., Xue, M., He, M., Huang, Z.J., and Scanziani, M. (2013). Inhibition of inhibition in visual cortex: the logic of connections between molecularly distinct interneurons. Nat Neurosci 16, 1068–1076.

Pluta, S., Naka, A., Veit, J., Telian, G., Yao, L., Hakim, R., Taylor, D., and Adesnik, H. (2015). A direct translaminar inhibitory circuit tunes cortical output. Nat Neurosci 18, 1631–1640.

Poort, J., Khan, A.G., Pachitariu, M., Nemri, A., Orsolic, I., Krupic, J., Bauza, M., Sahani, M., Keller, G.B., Mrsic-Flogel, T.D., and Hofer, S.B. (2015). Learning Enhances Sensory and Multiple Non-sensory Representations in Primary Visual Cortex. Neuron 86, 1478–1490.

Poort, J., Self, M.W., van Vugt, B., Malkki, H., and Roelfsema, P.R. (2016). Texture Segregation Causes Early Figure Enhancement and Later Ground Suppression in Areas V1 and V4 of Visual Cortex. Cereb Cortex 26, 3964–3976.

Reynolds, J.H., and Chelazzi, L. (2004). Attentional modulation of visual processing. Annu Rev Neurosci 27, 611–647.

Rudy, B., Fishell, G., Lee, S., and Hjerling-Leffler, J. (2011). Three groups of interneurons account for nearly 100% of neocortical GABAergic neurons. Dev Neurobiol 71, 45–61.

Sachidhanandam, S., Sreenivasan, V., Kyriakatos, A., Kremer, Y., and Petersen, C.C. (2013). Membrane potential correlates of sensory perception in mouse barrel cortex. Nat Neurosci 16, 1671–1677.

Saleem, A.B., Ayaz, A., Jeffery, K.J., Harris, K.D., and Carandini, M. (2013). Integration of visual motion and locomotion in mouse visual cortex. Nat Neurosci 16, 1864–1869.

Saleem, A.B., Lien, A.D., Krumin, M., Haider, B., Roson, M.R., Ayaz, A., Reinhold, K., Busse, L., Carandini, M., and Harris, K.D. (2017). Subcortical Source and Modulation of the Narrowband Gamma Oscillation in Mouse Visual Cortex. Neuron 93, 315–322.

Smith, M.A., Jia, X., Zandvakili, A., and Kohn, A. (2013). Laminar dependence of neuronal correlations in visual cortex. J Neurophysiol 109, 940–947.

Snyder, A.C., Morais, M.J., and Smith, M.A. (2016). Dynamics of excitatory and inhibitory networks are differentially altered by selective attention. J Neurophysiol 116, 1807–1820.

Soares, D., Goldrick, I., Lemon, R.N., Kraskov, A., Greensmith, L., and Kalmar, B. (2017). Expression of Kv3.1b potassium channel is widespread in macaque motor cortex pyramidal cells: A histological comparison between rat and macaque. J Comp Neurol 525, 2164–2174.

Spitzer, H., Desimone, R., and Moran, J. (1988). Increased attention enhances both behavioral and neuronal performance. Science 240, 338–340.

Vaiceliunaite, A., Erisken, S., Franzen, F., Katzner, S., and Busse, L. (2013). Spatial integration in mouse primary visual cortex. J Neurophysiol 110, 964–972.

Vigneswaran, G., Kraskov, A., and Lemon, R.N. (2011). Large identified pyramidal cells in macaque motor and premotor cortex exhibit thin spikes: implications for cell type classification. J Neurosci 31, 14235–14242.

Vinck, M., Batista-Brito, R., Knoblich, U., and Cardin, J.A. (2015). Arousal and locomotion make distinct contributions to cortical activity patterns and visual encoding. Neuron 86, 740–754.

Womelsdorf, T., Fries, P., Mitra, P.P., and Desimone, R. (2006). Gamma-band synchronization in visual cortex predicts speed of change detection. Nature 439, 733–736.

